# Underlying driving forces of the SARS-CoV-2 evolution: immune evasion and ACE2 binding affinity

**DOI:** 10.1101/2023.02.06.527236

**Authors:** Wentai Ma, Haoyi Fu, Fanchong Jian, Yunlong Cao, Mingkun Li

## Abstract

The evolution of SARS-CoV-2 is characterized by the emergence of new variants with a sheer number of mutations compared to their predecessors, which conferred resistance to pre-existing antibodies and/or increased transmissibility. The recently emerged Omicron subvariants also exhibit a strong tendency for immune evasion, suggesting adaptive evolution. However, previous studies have been limited to specific lineages or subsets of mutations, the overall evolutionary trajectory of SARS-CoV-2 and the underlying driving forces are still not fully understood. In this study, we analyzed the mutations present in all open-access SARS-CoV-2 genomes (until November 2022) and correlated the mutation’s incidence and fitness change with its impact on immune evasion and ACE2 binding affinity. Our results showed that the Omicron lineage had an accelerated mutation rate in the RBD region, while the mutation incidence in other genomic regions did not change dramatically over time. Moreover, mutations in the RBD region (but not in any other genomic regions) exhibited a lineage-specific pattern and tended to become more aggregated over time, and the mutation incidence was positively correlated with the strength of antibody pressure on the specific position. Additionally, the incidence of mutation was also positively correlated with changes in ACE2 binding affinity, but with a lower correlation coefficient than with immune evasion. In contrast, the mutation’s effect on fitness was more closely correlated with changes in ACE2 binding affinity than immune evasion. In conclusion, our results suggest that immune evasion and ACE2 binding affinity play significant and diverse roles in the evolution of SARS-CoV-2.

## Introduction

Recent SARS-CoV-2 lineages (e.g., BA.2 sub-lineages, BA.5 sub-lineages) have an average of over 80 mutations relative to the earliest genome (www.nextstrain.org). Some variants exhibited significant alterations in their transmissibility, antigenicity, and pathogenicity compared to their predecessors^1-3^. The most notable ones are those referred to as Variants of Concern (VOC), including Alpha, Beta, Delta, Gamma, and Omicron, which are more transmissible or/and able to escape the pre-existing immune pressures, and thus led to multiple surges of infection peaks on a local or global scale. Intrinsic transmissibility and immune pressure have been proposed as the two primary forces driving the evolution of SARS-CoV-2 as well as other viruses^4^. For example, the rapidly spreading D614G and N501Y mutations could enhance viral transmission by increasing the ACE2 binding affinity^5,6^, while the E484K reduces susceptibility to neutralizing antibodies^7,8^. Moreover, with the development and optimization of the high-throughput deep mutational scanning (DMS) method, it became more feasible to access the effect of RBD mutations in the binding affinity to antibody^9,10^ and human ACE2 receptor^11,12^, which facilitated the identification of a number of RBD mutations in Omicron and its sub-lineages that conferred significant immune evasion against antibodies induced by prior infections or vaccinations, while maintaining sufficient binding affinity to the human ACE2^11,13^. Moreover, recent studies proposed an unprecedented convergent evolution of BA.2 and BA.4/5 subvariants (e.g., BQ.1, XBB, BM.1), which enable a near-complete evasion against most known antibodies, emphasizing the significant role of immune pressure on SARS-CoV-2 evolution^14,15^. However, previous studies mainly focused on a few fast-growing variants and mutations that were known to have a great impact on immune evasion, and thus may suffer from survival bias, i.e., the findings may not be applicable to other variants that account for a large fraction of the data. Therefore, a thorough investigation of all mutations, including those not define any virus lineage and are found in a small number of samples, is needed to disentangle the evolutionary trajectory and underlying evolutionary driving forces of SARS-CoV-2.

Besides, there are also other unanswered scientific questions concerning the evolution of SARS-CoV-2. For example, 1) Did adaptive evolution only occur in the RBD region, which is the primary target of neutralizing antibodies and to which human ACE2 binds? 2) Did the SARS-CoV-2 evolutionary pattern vary over time and between different lineages? 3) Whether the two factors—immune pressure and intrinsic transmissibility—contribute equally to viral evolution and do their effects change over time? The evolution of SARS-CoV-2 would be better understood if the above questions could be answered.

In this study, we have investigated the mutations inferred from more than six million open-access SARS-CoV-2 sequences as of Nov 23, 2022, and correlated the mutation spectrum and incidence with their immune-evasive and ACE2 binding potentials estimated from the DMS and neutralization data. We found that different SARS-CoV-2 macro-lineages exhibited distinct mutation patterns in the RBD region that are likely to be associated with continually changing humoral immune pressures, and the immune pressure-driven mutation became more evident in the recent BA.2 and BA.4/5 sub-lineages. Besides, the ACE2 binding affinity also played a significant role in the viral evolution, especially at the early stage after the emergence of new variants when the humoral immune pressure was relatively low. Although the immune pressure was more correlated with the occurrence of mutations compared to the ACE2 binding affinity, it is interesting to note that enhanced ACE2 binding affinity was more closely correlated with increased virus fitness than immune evasion.

## Results

### The accelerated mutation rate in the SARS-CoV-2 RBD region

Recently, a large number of immune-evasive BA.2 and BA.4/5 subvariants were emerging in the population^15,16^, implying the evolution of SARS-CoV-2 might be accelerating. To verify this hypothesis, we retrieved 200 sequences per month between January 2020 and November 2022 to estimate the mutation rate of SARS-CoV-2 over time, the automatic piecewise linear regression analysis found two turning points associated with a dramatic rise in the mutation number (Figure 1A), corresponding to the emergence of the Alpha and the Omicron variants, which had a sheer number of mutations relative to their possible predecessors. The slope of the regression lines before and after the turning points was similar, indicating that the mutation rate did not significantly increase over time. Meanwhile, the analysis performed on different VOCs confirmed that the mutation rate did not increase in the latest Omicron sub-lineages (Figure 1B). However, we found that the Omicron variants (BA.1, BA.2, BA.4/5) showed an accelerated rate of amino acid changes compared to previous viral lineages in the RBD region (3-83 folds higher) (Figure1C).

**Figure 1.**
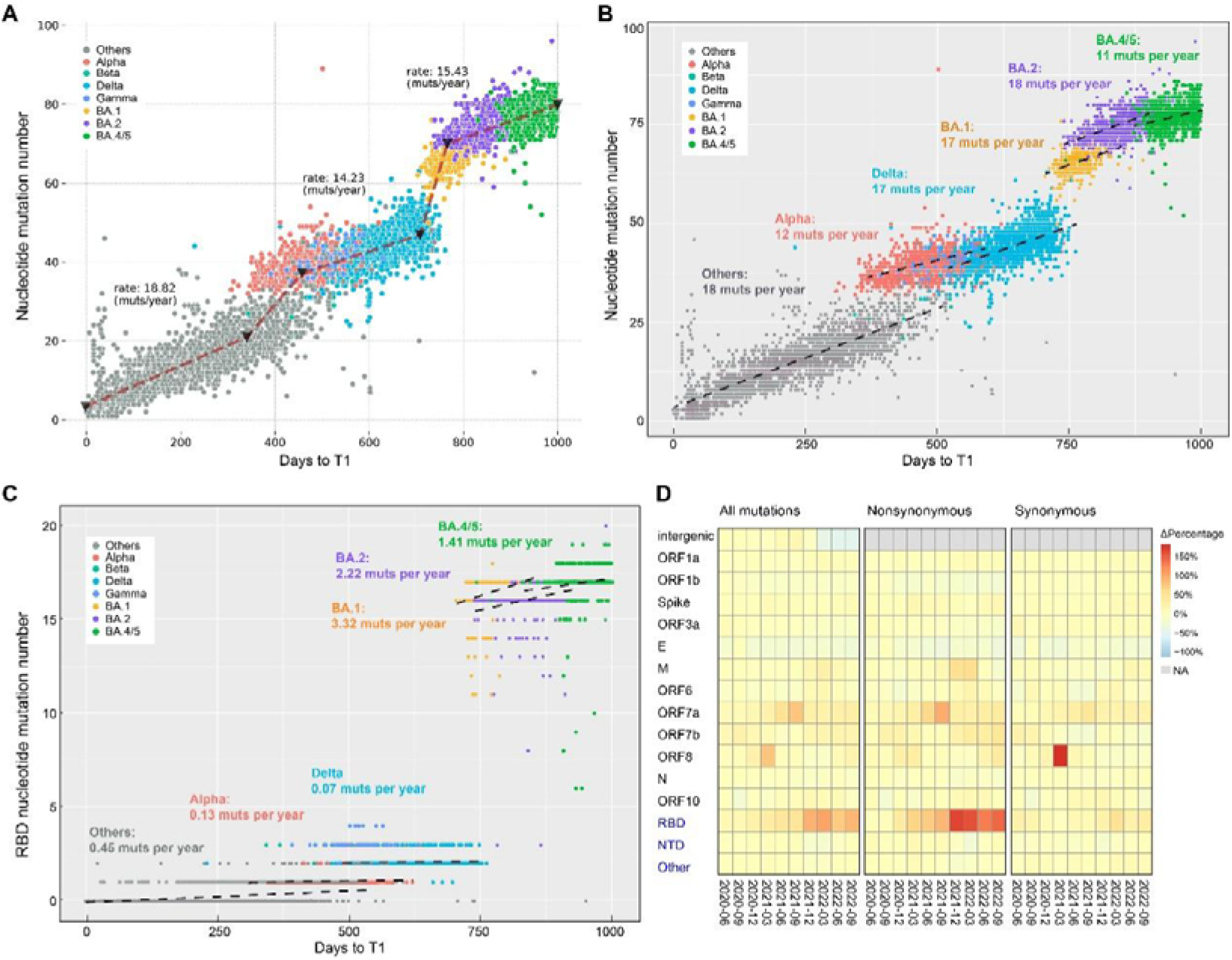
The mutations in the SARS-CoV-2 genome. A) The correlation between mutation count and collection date. Two hundred sequences were randomly selected from each month based on the collection date. The segmented regression line was fitted using automatic piecewise linear regression, and the mutation rate was estimated as the slope of the regression line. T1 represents Dec 24, 2019, which is the collection date of the first open-access SARS-CoV-2 sequence. B) The correlation between mutation count and collection date for major lineages. The estimated mutation rates for six major lineages are shown on top of the linear regression lines. C) The mutation rate of the RBD region (residues 331-531 of the *Spike* gene) in major lineages. D) The distribution of mutations across different genomic regions over time. The three regions (RBD, NTD, and other regions) in the *Spike* gene are separately shown in the figures (marked in blue). The time window size is three months, with the first three months (March, April, and May in 2020) used as reference (samples collected prior to March are limited and were not included in the analysis). The cell’s color indicates the degree of change relative to the reference.

To further verify the accelerated mutation rate in the RBD region and test whether other genes were also subject to a higher mutation rate in recent Omicron sublineages, we retrieved all mutations inferred from the UShER mutation-annotated-tree constructed from 6,484,070 complete high-quality SARS-CoV-2 genomes^17^. We found that the proportion of mutations in the RBD region increased over time, which was mainly contributed by the excess number of non-synonymous mutations (Figure 1D), i.e., the proportion of non-synonymous mutations in the RBD regions increased from 52.8% in the early stage of the pandemic to 62.1% in the recent Omicron sub-lineages. However, the same tendency was not observed in other genes or other regions in the *Spike* gene.

### RBD Mutations showed a lineage-specific pattern and became more aggregated in recent lineages

To further investigate the evolution of the sequences in the RBD region, we analyzed the distribution and incidence of all amino acid mutations identified in this region, which included 855 mutations that accounted for 6,328 mutation events. First, we found that the incidence differed among different mutations. The overall P50 (proportion of mutations that accounted for 50% of all mutations events) was 7.5%, with the proportion decreasing over time (Figure 2A), suggesting the mutation tended to become more aggregated in recent lineages. Second, the high-frequency mutations tended to be shared among variants belonging to the same macro-lineages. The viral lineages (with ≥ 6,000 sequences) could be classified into four clusters according to the similarity of the incidence of 59 mutations that showed the highest incidences (top five) in at least one lineage (Figure 2BC). It is interesting to note that these four clusters correspond to four viral macro-lineages (B.1, Delta, BA.1/2, and BA.4/5), suggesting that the variants belonging to the same lineages, including those circulated at the same time, tended to have the same high-frequency mutations, which is a signature of convergent evolution. Third, mutations occurring at the same position could be of different types. Forty high-frequency mutations were found at 16 positions (Figure S1), while the other 19 high-frequency mutations were found at 19 distinct positions. Different high-frequency mutations at the same position differed in amino acid polarity, acidity, and charge, thus may result in varying effects on the affinity between the virus and the host cell and antibodies. Meanwhile, we also noted that distinct mutations at the same position could arise in the variants belonging to the same lineage, implying that the virus may react differently to the same pressures.

**Figure 2.**
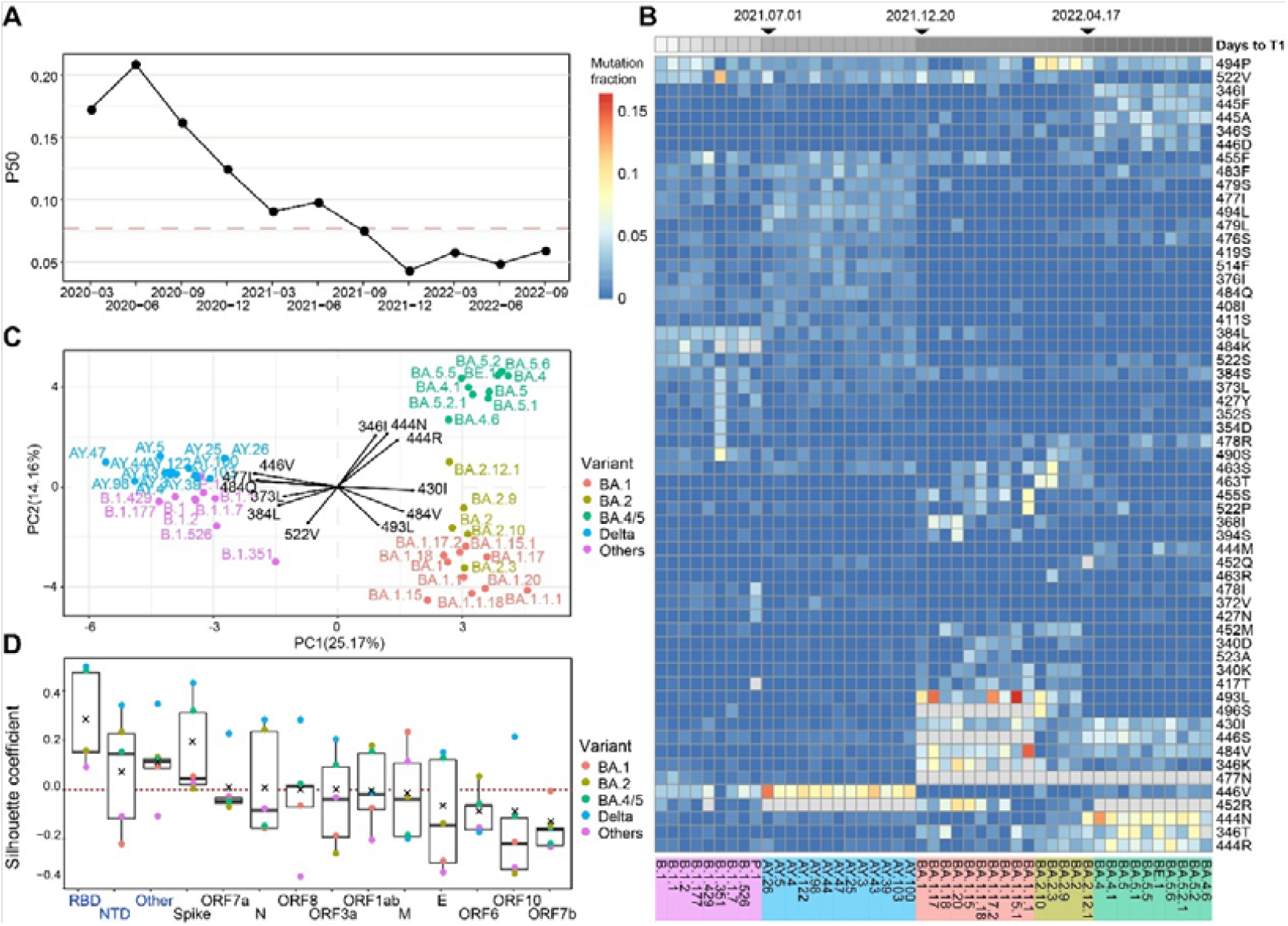
The convergent evolution of the RBD sequences in the SARS-CoV-2 genome. A) The proportion of mutations that accounted for 50% of all mutations events (P50) at different time points. The red dashed line indicated the P50 for all mutations. B) The mutation incidence in various SARS-CoV-2 lineages. The color denoted the ratio of this mutation’s frequency to the frequency of all mutations in the lineage. The top five most frequent mutations in each lineage are shown. Mutations that had been fixed in the lineage are labeled in grey. C) The Principal component analysis (PCA) plot of mutation incidence across different lineages. The input data were taken from Figure 2B. The top three mutations that explained the highest variance in each quadrant are labeled in the figure. D) The clustering significance of different lineages based on mutation incidence. The silhouette coefficient of the clustering of lineages in various genomic regions is shown. The analysis included 40 lineages with over 6,000 sequences. Lineages are sorted by the collection date of the earliest 5% of sequences belonging to each lineage. To prevent errors caused by misplaced sequences in the phylogenetic tree, back/reverse mutations were excluded from the analysis.

The high-frequency mutations in B.1 and Delta clusters were similar, which featured by the high incidence of 373L, 384L, 552V, 484Q, 477I, and 446V mutations compared to the Omicron clusters, with the former three more frequently observed in the B.1 cluster, and the latter three were more frequently observed in the Delta cluster. Particularly, the mutation 446V was 2.1 times more frequent in the Delta cluster compared to that in the B.1 cluster. The difference between the BA.1/2 and BA.4/5 clusters was more remarkable compared to that between B.1 and Delta clusters, despite that some of the BA.2 and BA.4/5 subvariants were circulating at the same time in the population. BA.1/2 had a higher incidence of 484V and 493L, while BA.4/5 had a higher incidence of 444R/N and 346I/T.

To test whether the convergent mutation also occurred in other genomic regions, the same analysis was conducted in all genes on the SARS-CoV-2 genome. We found that the same trend was not observed in other genes, as indicated by an average Silhouette value less than 0, which denotes that the similarity of the mutation pattern within the same lineages is lower than that between different lineages (Figure 2D). Noteworthy, the NTD region and other regions in the *Spike* gene showed signs of convergent mutations in some lineages, but the magnitude is much lower than that in the RBD region.

### Humoral immune pressure is correlated with the evolution of the SARS-CoV-2

To investigate whether the convergent evolution in the RBD region can be explained by a similar humoral immune pressure from antibodies acting on the same macro-lineage, we classified the mutations into immune escape mutations and non-immune escape mutations according to the DMS scores against over 2,000 antibodies that belong to 12 epitope groups (hereafter referred to as “antibody types”). The mutation that significantly reduced the affinity to any of the 12 antibody types was defined as an immune escape mutation. First, we noted that the proportion of immune escape mutations was higher in the Omicron lineages than in other earlier lineages (47.3% *vs*. 67.0%, pvalue<0.001). Meanwhile, the spectrum of immune evasion caused by mutations changed markedly over time and tended to be more concentrated on specific antibody types in recent lineages (Figure 3A). Specifically, mutations escaping D1 and D2 antibody types became more enriched in the Delta lineages compared to the earlier lineages; Omicron BA.1/2 sub-lineages exhibited an increased proportion of mutations escaping A, C, E2.1, and E2.2 antibodies; and Omicron BA.4/5 sub-lineages showed the strongest immune escape against D1 and D2 antibody types.

**Figure 3.**
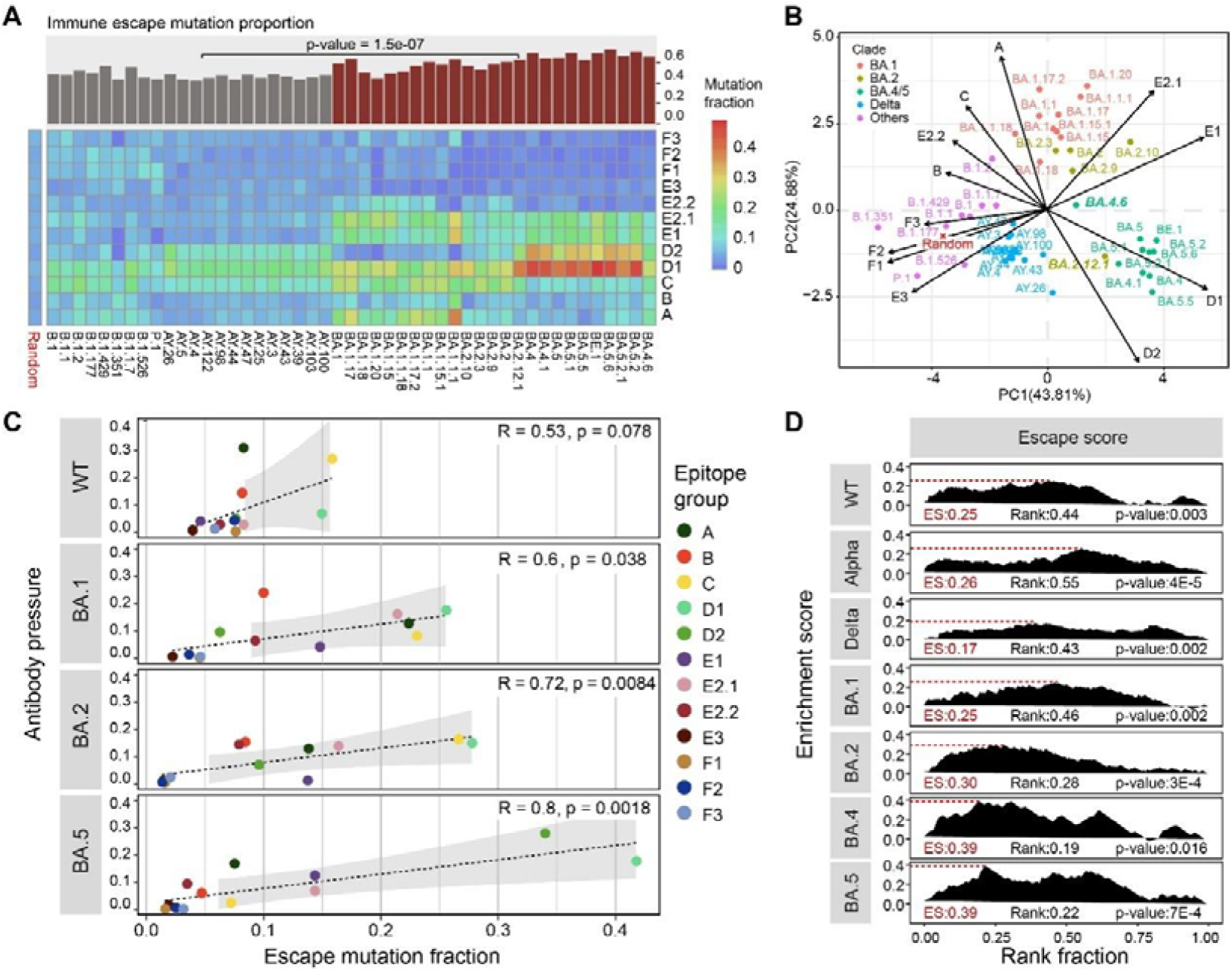
The correlation between the incidence of RBD mutations and humoral immune pressure. A) The distribution of escape mutations in 12 antibody epitope groups. The bar plot on top shows the proportion of immune escape mutations among all mutation events in different lineages. The null distribution of mutations, assuming no epitope group preference, is shown on the left side of the heatmap. B) PCA plot of the mutation distribution in different lineages. The input data were taken from Figure 3A. C) The correlation between the prevalence of escape mutations and antibody pressure in twelve epitope regions. The x-axis represents the proportion of immune escape mutations in a particular epitope region among all escape mutations. The y-axis represents the proportion of immune pressure exerted on a particular epitope region, calculated by summing the neutralizing activities of all antibodies that belong to this epitope group. D) The correlation between immune evasion capacity and mutation incidence. All mutations in each major lineage were reverse-sorted by their incidences, and gene set enrichment analysis (GSEA) was conducted to examine whether the high-weight escape mutations were enriched among the high-incidence mutations (details are provided in the Methods section). The ES value represents the highest cumulative score (the peak), the Rank value represents the x-axis position of the peak, and the p-value was calculated through 1,000 runs of random reordering. WT stands for wide type, which has no mutation compared to the NC_045512.2 in the RBD region.

We also noted that the variants with similar background RBD sequences (belonging to the same macro-lineage) tended to have the mutations that evaded the same antibody types (Figure 3B), suggesting that the immune pressure was similar among variants in the same macro-lineage. Meanwhile, we also found that the immune pressure was altered with the accumulation of immune escape mutations. For example, the variant BA.2.12.1 acquired an extra 452Q mutations compared to its predecessor BA.2 variant, this mutation could facilitate the escape from the E2 and D1 antibody types according to the DMS data, making its antigenicity more similar to BA.4/5, which had a 452R mutation that could evade the same antibody types. Accordingly, the mutation pattern of BA.2.12.1 is more similar to BA.4/5 subvariants than other BA.2 subvariants. Additionally, the variant BA.4.6 possessed an additional RBD mutation 346T compared to other BA.4 lineages, which is capable of compromising the efficacy of D1, E1, and E2.1 antibody types, hence displayed a distinctive pattern of immune-evasive mutations compared to other BA.4/5 (Figure 3AB).

To further quantify the impact of humoral immune pressure on the immune-evasive mutations in the RBD region, we estimated the immune pressure on four major variants (WT, BA.1, BA.2, and BA.4/5) from each antibody type using the pseudovirus-neutralization data (antibodies induced by different variants were retrieved from previous studies)^13,14^. A marginal correlation was found between the incidence of the immune escape mutation and the immune pressure on the WT variants (Figure 3C). The correlation became more significant in the Omicron lineages and the correlation coefficients increased over time, suggesting a stronger immune pressure on recent lineages, which may be related to the increased antibody prevalence in the population as a result of mass vaccination campaigns and infections. Moreover, Gene Set Enrichment Analysis (GSEA) indicated that the high-frequency mutations were more likely to confer a stronger resistance to the highly potent antibodies against the variant (Figure 3D), and the tendency was more remarkable in the Omicron lineages, implying that the humoral immune pressure played a stronger role in the recent evolution of SARS-CoV-2.

### Increased ACE2 binding affinity may also drive the evolution of SARS-CoV-2

Besides immune escape, increased transmissibility is another major direction of viral evolution^4,18^. One of the variables that correlated with viral transmissibility is ACE2 binding affinity, which determines how easily an infection/transmission is established. By analyzing the effect of the mutation on the ACE2 binding affinity under different genetic backgrounds (WT, Alpha, Delta, BA.1, BA.2, BA.4/5), which was measured by high-throughput DMS screening^12^, we found that 34% of the mutations that occurred in the WT had the potential to enhance ACE2 binding affinity, while the proportion decreased to 19% and 23% in the Alpha and Delta lineages, respectively (Figure 4A). In the Omicron lineages, the proportion was 32% and 33% in the BA.1 and BA.2 sub-lineages, respectively, and it dropped to 18% in the BA.4/5 sub-lineages. Meanwhile, the proportion of immune escape mutations also varied amongst different macro-lineages, with Omicron lineages having a higher proportion of immune escape mutations compared to earlier variants. The dynamic of the mutation pattern may reflect a shift in the force that drove the viral evolution.

**Figure 4.**
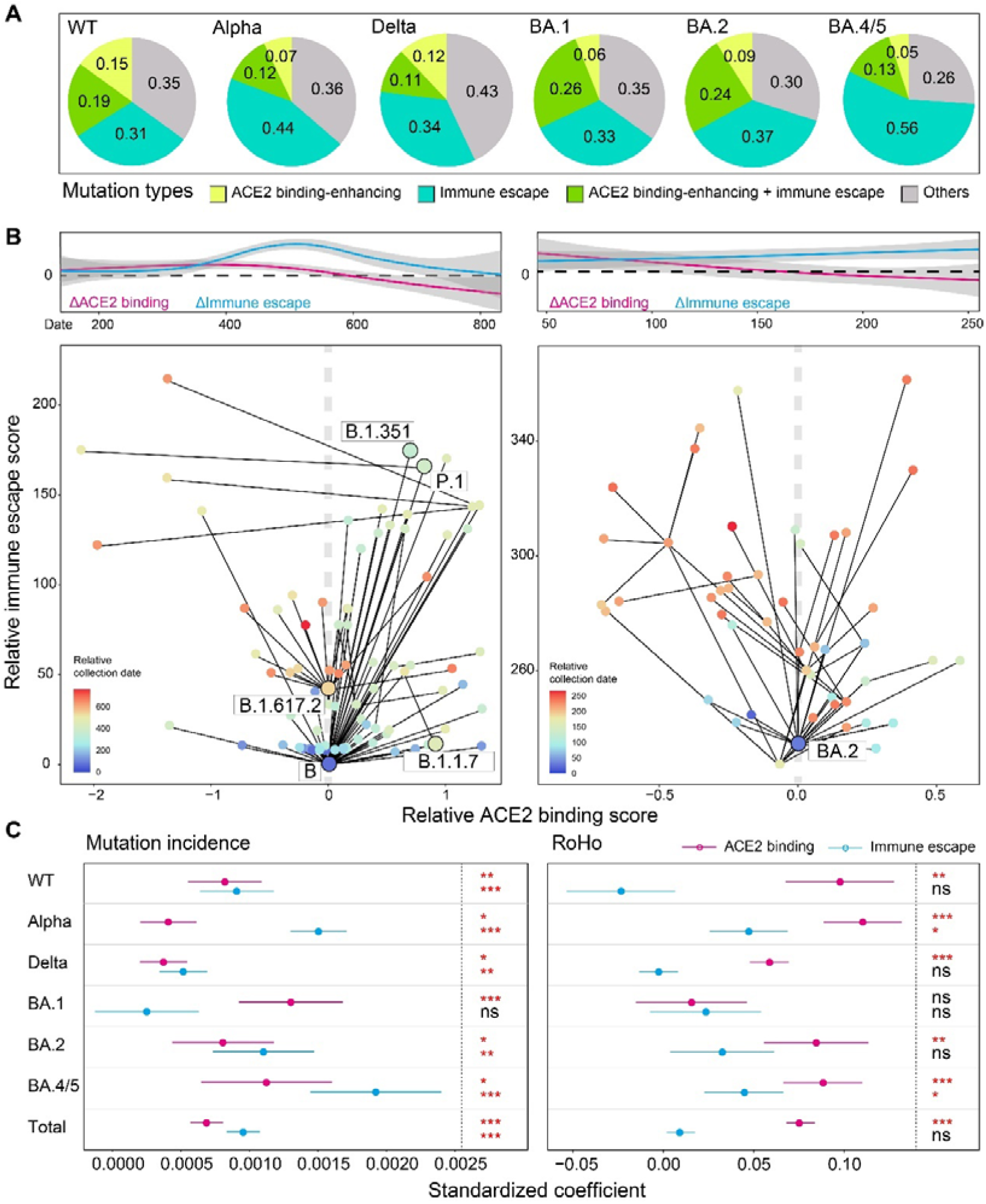
Immune evasion and ACE2 binding affinity drove the evolution of SARS-CoV-2. A) The composition of different mutation types in six major lineages. The ACE2 binding-enhancing mutations refer to those having a positive ACE2 binding score (the sum of ACE2 binding score and RBD expression score). The immune escape mutations refer to those having an escape score that is three times greater than the average escape score of all mutations in at least one antibody epitope group. B) The mutation trajectory of two major variants B and BA.2. The relative ACE2 score and immune escape score of each variant represent the change in comparison to the ancestor variant. The regression lines of the two variables were obtained using a generalized additive model and are shown on top of the figures. C) The correlation between mutation incidence (left) / fitness change (right) and the ACE2 binding and immune escape scores of the mutation. The correlation coefficient and significance were obtained from a multivariate linear regression. * p<0.05, ** p<0.01, *** p<0.001, ns: not significant.

When tracing the evolution of the two major variants B and BA.2 that spread over a long period of time, we found that mutations that occurred in the early stage in the two lineages were more likely to enhance the ACE2 binding affinity than that occurred in the later stage, suggesting that increased binding affinity was more advantageous for the viral transmission when a new variant emerged in the population. In contrast, later-stage mutations were more likely to facilitate immune evasion compared to the early-stage mutations, suggesting that immune evasion is more advantageous for viral transmission when a large population has been infected by the variant or herd immunity had been established.

Through multiple linear regression analysis, we found that both immune evasion and ACE2 binding affinity independently correlated with the mutation incidence in all macro-lineages, with the former having a greater impact on the mutation incidence in all lineages except the BA.1. Notably, immune evasion showed the highest standardized correlation coefficients in the BA.4/5 sub-lineages, which is consistent with the observation that the recent Omicron subvariants showed an accelerated rate to evade the antibody induced by earlier variants infections^14^. However, the preponderance of specific mutation types during particular pandemic stages did not necessarily mean that these mutations were advantageous to the virus’s fitness. To evaluate the effect of mutation on the fitness of the new variant, we calculated the RoHo score for each pair of new variants and its predecessor^19^, which differed by one or two mutations, to represent the fitness advantage of the mutation. We found that the ACE2 binding affinity of the mutation showed a more significant correlation with the fitness advantage than immune evasion, while the contribution of immune evasion to the variant fitness was greater in the recent BA.4/5 subvariants.

## Discussion

In the study, we have provided the first comprehensive evolutionary analysis of SARS-CoV-2 from the perspective of immune evasion and ACE2 binding affinity properties, which were the only two functional features available for a large number of mutations. Although other factors, such as the virus particle stability, replication efficiency, incubation time, and cell tropism may also be involved in the evolution of SARS-CoV-2, the lack of relevant data prevented us from taking them into account in our study^20^.

We found that the mutation rate of SARS-CoV-2 was similar in different macro-lineages. Estimation of the mutation rate in other studies was much higher than the segmented mutation rates estimated in our study^21,22^, because the emergence of the variants that had a sheer number of mutations (e.g., Alpha, Omicron) compared to the then circulating variants was not considered our estimation. As the underlying mechanism for the emergence of these new VOCs is still mysterious^23,24^, our study only focused on how the variant evolved in the population after their emergence. The distribution of mutations on the viral genome changed over time, with an increase in the proportion of mutations in the RBD region since early 2021. Notably, the rise is primarily attributable to an increase in non-synonymous mutations, suggesting that the RBD region is under growing positive selection. Given that the RBD region is where the most neutralizing antibodies and host cells bind to^25^, mutations in this region could have a significant impact on the virus’s ability to evade the immune system and spread to other cells, thus it is not surprising that this region would be subject to stronger natural selection. The increased selection pressure on the RBD region over time might be caused by the rising vaccination and infection rates, which resulted in highly concentrated humoral immune pressure.

Convergent evolution has been observed in recent Omicron subvariants, and presumably caused by the concentrated humoral immunity pressures^14^. Our study indicated that the convergent evolution was present even at the beginning of the pandemic, albeit to a lesser extent. The low-intensity convergent evolution may reflect mutation bias or low levels of immunity pressure. Besides the immune escape mutations (446V, 484Q, 346I, 444N/R, 430I, 494P, 484V) and ACE2 binding-enhancing mutations (384L), there were other convergent mutations whose functions are not known (477I, 373L, 522V) repeatedly occurred in particular lineages, the underlying mechanism required further investigation. In this study, only the RBD region showed a significant convergent evolution among high-frequency mutations, however, this did not rule out the possibility that convergent mutation will take place in other regions in the future when the predominant immune pressure switches to other regions. For example, Y144 deletion has been frequently observed in the Omicron and early variants^26^, which confers resistance to the neutralizing antibody targeting the NTD region, other mutations may arise in this region when most RBD-targeting antibodies were escaped.

We noted that the most prevalent convergent mutations included distinct mutation types occurring at the same position, suggesting that diverse mutation types may be able to offset the same selection pressure. However, the mutation type could also be lineage-specific at some positions, suggesting that different mutation types at the same location may have distinct functional effects. For example, 346K is primarily observed in the BA.1 subvariants, while 346T/S/I was more abundant in the BA.4/5 subvariants^2^. We hypothesize that the above phenomenon may reflect the shifting pressures on the virus at different phases. When Omicron first emerged in the population, it has an overwhelming growth advantage over Delta variants due to the numerous immune-evasive mutations in the Omicron RBD region. The high prevalence of 346K in BA.1 might be explained by the higher mutation rate from A to G (resulting in 346K) than the mutations from A to C/T (346S) and mutation from G to C (346T) in the SARS-CoV-2 genome^27^. The immune pressure against the virus became much stronger after Omicron infected a large proportion of the population, hence immune escape mutations offered a higher transmission advantage than the ACE2-binding enhancing mutations at the later stage. Then, mutation 346T, which offers the highest immune evasion under the BA.4/5 genetic background (Table S1), became the most predominant mutation in the BA.4/5 subvariants. This hypothesis coincides with our finding that the immune escape mutations were more common once the variant infected a large proportion of the population, while the ACE2 binding-enhancing mutations were more prevalent when a new variant (with a significant antigenicity change) first appeared (Figure 4).

We found that the occurrence of new mutations was significantly correlated with the immune evasion as well as ACE2 binding affinity, agreeing with previous studies based on limited lineages and mutations^3^, and thus confirmed a function-driven virus evolution. Moreover, we have quantified the effects of the two factors on mutation occurrence and mutation fitness under different genomic backgrounds, whose effects are difficult to distinguish using incomplete data, as the same mutation occurred multiple times in different lineages, where their impact may vary. Overall, the immune evasion showed a stronger correlation with mutation incidence than increased ACE2 binding affinity. Given that RNA virus has a high mutation rate, the large number of viruses in the human body could produce a population of viruses with a great genetic diversity^28-30^. The viral population would then be subject to selection by antibodies induced by infections, and the variants with mutations conferring higher resistance to antibodies would have a superior replication advantage over other variants, making them more likely to dominate the viral population and be observed as a mutation at the individual level. Meanwhile, we found that the correlation coefficient between immune evasion and mutation occurrence increased remarkably in the recent Omicron subvariants of BA.5, which is consistent with the recent observation that a large number of immune evasive mutations were found in Omicron subvariants^15^. This tendency is likely owing to the imprinted humoral immunity, which resulted in reduced diversity of neutralizing antibodies and thus exerted more concentrated immunological pressure on recent variants^14,31^.

Compared to the immune evasion, ACE2 binding affinity exhibited a less significant correlation with the mutation occurrence, but a more significant correlation with viral fitness. Although previous studies have reported that ACE2 binding-enhancing mutations can promote viral transmission^1,32^, including the recent acquisition of S486P in XBB.1.5, which could significantly increase the ACE2 binding affinity^33^, the importance of ACE2 binding affinity is overlooked in previous studies of SARS-CoV-2 evolution. We hypothesize that the impact of increased ACE2 binding affinity on viral fitness is more universal compared to immune evasion, while the effect of the latter is more context-dependent. Namely, all mutations that increase the ACE2 binding effect confer a transmission advantage as an infection is much easier to be established^34^, whereas the immune-evasive mutations provide additional transmission advantage only in the population that has been infected by the prototype (probably also needs to be in the recent past when the neutralizing antibody titer is high). Therefore, as the rate of reinfection rises in the Omicron era, we would expect that immune evasion will contribute more to the viral transmission advantage in the future.

Our study has several limitations. First, the effects of mutations on immune evasion and ACE2 binding affinity were estimated based on the data generated on limited RBD backgrounds, thus the interaction between mutations, as demonstrated in previous studies^12,35^, was not considered when there are multiple mutations relative to the background sequences. Second, while both the humoral and cellular immune systems exert pressure on the virus, only the former was taken into account in this study as data on the latter is not yet available. Third, the antibody compositions were estimated from a small number of samples infected by a specific variant, thus may not accurately reflect the humoral immune pressure on the virus due to the complex history of vaccination and infection across different populations. Additionally, the quantity of the antibodies in the human body is unknown due to technical limitations, and therefore, different antibodies were quantitatively equally weighted when estimating the immune pressure on the virus, potentially leading to biased estimates.

The unprecedented number of SARS-CoV-2 viral genomes enables us to track the trajectory of SARS-CoV-2 evolution. Meanwhile, advancements in technologies and the accumulated data have greatly advanced our understanding of the mutation function. Our study showed a significant correlation between the immune evasion and ACE2 binding affinity of the mutation and mutation incidence and fitness change, which improved our understanding of the underlying forces driving the evolution of SARS-CoV-2. However, we are still far from being able to precisely predict the direction of viral evolution due to a lack of comprehensive functional data on mutations and their interactions.

## Methods

### Estimation of the mutation rate of SARS-CoV-2

A total of 6,484,070 high-quality open-access SARS-CoV-2 sequences and corresponding metadata were downloaded from the UShER website on November 23^rd^, 2022^17^. To calculate the mutation rate, 200 sequences from each month were randomly selected based on their collection date. The number of mutations (relative to the NC_045512.2) was analyzed using the Optimal Piecewise Linear Regression Analysis (OPLRA) method, a mathematical programming technique that divides the data into optimal segments and fits a linear regression function to each segment to minimize the overall absolute error^36^. The Akaike and the Bayesian metrics were employed to balance predictive accuracy and model complexity to determine the optimal number of segments^37^. Meanwhile, the mutation rate was also estimated for six macro-lineages (WT, Alpha, Delta, Omicron BA.1, BA.2, and BA.4/5) using a linear regression model. In addition, The RoHo value of the mutation, which is the ratio of the number of descendants in sister clades with and without a specific mutation^19^, was obtained using the matUtils tool from the UShER toolkit to represent its fitness.

### Estimation of the mutation incidence from the UShER phylogenetic tree

The mutation events were retrieved from the masked UShER mutation-annotated tree. First, the matUtils tool from the UShER toolkit was used to convert the protocol buffer format to the JSON format. Then, to reduce the number of false positive events caused by the incorrect placement of the sequence in the phylogenetic tree, which included an unprecedented number of sequences, a mutation event was called only from leaf nodes (i.e., real sequences) or internal nodes that have at least one offspring being a leaf node. Meanwhile, no more than two mutations were allowed between those nodes and their parental nodes. The number of mutation events identified on the phylogenetic tree was used to represent the incidence of mutation. Only viral lineages with more than 6,000 sequences were considered for comparing the mutation incidence between different lineages to minimize the bias caused by a small sample size.

### Estimation of the mutation escape score and humoral immune pressure

The antibody spectrum, neutralizing activity, antibody epitope group, and mutation escape score were obtained from a previous study^14^. Briefly, 2,170 antibodies were identified from the sera of vaccinated individuals and convalescent patients of the wild type (WT), BA.1, BA.2, and BA.5 variants using single-cell V(D)J sequencing. The neutralizing activities of these antibodies against the WT, BA.1, BA.2, and BA.5 variants were determined using a pseudovirus neutralization assay. The impact of single amino acid mutation in the RBD region on the neutralization efficacy of antibodies was assessed using a high-throughput deep mutation scanning strategy. For each mutation, an escape score was computed by fitting an epistasis model to reflect the degree of the change in antibody neutralization capacity caused by the mutation^38,39^. The raw escape score for each antibody was normalized by the max score among all mutations.

The antibodies were classified into 12 epitope groups according to their mutational escape profiles based on the WT using multidimensional scaling (MDS) followed by k-means clustering^13,14^. Thus, the antibodies in the same epitope group tended to be escaped by the same set of mutations. For each epitope group, mutations with an escape score (the average of the scores against all antibodies belonging to the epitope group) that was three times higher than the average escape score of all mutations were defined as immune escape mutations. The immune pressure induced by each antibody epitope group was calculated by summing the neutralizing activity of all antibodies belonging to the group.

### Calculation of the correlation between immune-evasion capacity and mutation incidence

Mutations that can evade neutralizing antibodies can give the virus a transmission advantage. Gene set enrichment analysis (GSEA) was used to test the significance of the correlation between mutation incidence and the immune-evasion capacity of the mutation^40^. The immune-evasion capacity of the escape mutation for each antibody was calculated as the product of the escape score and the neutralizing activity of the antibody, and the values for all antibodies were summed up to represent the overall immune-evasion capacity of this mutation. The weight of the escape mutation (reward) was then set to be the proportion of immune evasion capacity contributed by the mutation relative to all mutations, while the weight of non-escape mutations (penalty) was set to be the reciprocal of the number of non-immune escape mutations, so the sum of the absolute weight of both mutation types was 1. All mutations in each lineage were reverse-sorted by incidence, and the GSEA was conducted to determine whether high-weight escape mutations were enriched among high-incidence mutations. The p-value indicating the significance of enrichment was calculated after 1,000 runs of random reordering. Notably, the escape score and the neutralizing activity for the WT variant were used for the Alpha and the Delta variants due to the lack of data for the latter two variants.

### Estimation of the ACE2 binding affinity

The DMS data for ACE2 binding and RBD expression were obtained from previous studies^12,41^, which included measurements on the WT, the Alpha, Beta, Delta, Omicron BA.1, and the BA.2 variants. Since ACE2 binding affinity and RBD expression are both critical for viral transmission, their effects were merged by summing their values (the two values are both log fold changes), a similar method was also used in a previous study^14^. The data for Omicron BA.2 was used to represent Omicron BA.4/5 due to the absence of data for BA.4/5.

## Acknowledgments

We thank all the scientists around the globe for performing SARS-CoV-2 sequencing and surveillance analysis. This study was funded by the National Natural Science Foundation of China (Grant No. 82161148009), the Strategic Priority Research Program of Chinese Academy of Sciences (Grant No. XDB38030400), and the Key Collaborative Research Program of the Alliance of International Science Organizations (ANSO-CR-KP-2022-09).

## Ethics statement

Not applicable.

## Author contributions

M.L. designed the study. W.M. and M.L. wrote the manuscript with input from all authors. W.M., H.F., and F.J. performed bioinformatics analyses. F.J. and Y.C. generated all the DMS and neutralization data.

