## Supplementary Figure 1 for "Underlying driving forces of the SARS-CoV-2 evolution: immune evasion and ACE2 binding affinity"

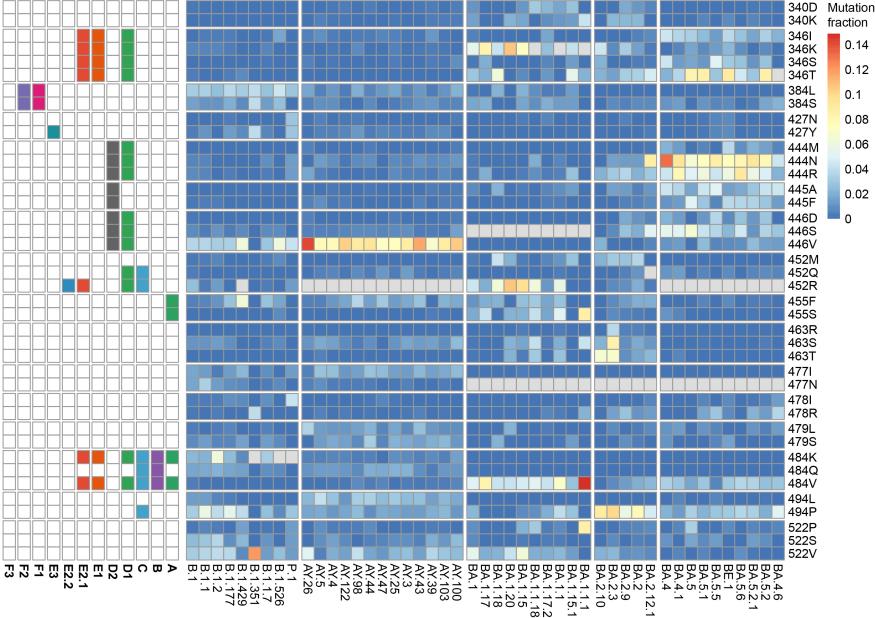

**Figure S1. The convergent evolution of the RBD region in the SARS-CoV-2 genome.** The color denoted the ratio of the mutation's frequency to the frequency of all mutations in the particular lineage. The top five mutations occurring most frequently in each lineage are shown, while the sites with just one high-frequency mutation were excluded. The mutations that had been fixed in the lineage were labeled in grey. The antibody epitope group that was evaded by the mutation in the right panel is illustrated in the left panel.
